# Arresting microbiome development limits immune system maturation and resistance to infection

**DOI:** 10.1101/2022.01.17.476513

**Authors:** Jean-Bernard Lubin, Jamal Green, Sarah Maddux, Lidiya Denu, Tereza Duranova, Matthew Lanza, Meghan Wynosky-Dolfi, Igor E. Brodsky, Paul J. Planet, Michael A. Silverman

## Abstract

Disruptions to the intestinal microbiome during weaning lead to long-term negative effects on host immune function. However, the critical host-microbe interactions occurring during weaning required for healthy immune system development remain poorly understood. We find that restricting microbiome maturation during weaning leads to stunted immune system development and increased susceptibility to enteric infection. We developed a gnotobiotic mouse model of the early-life microbiome designated as Pediatric Community (PedsCom). This nine-member consortium of microbes derived from intestinal microbiomes of preweaning mice stably colonized germfree adult mice and was efficiently transmitted to offspring for multiple generations. Unexpectedly, the relative abundance of PedsCom microbes were largely unaffected by the transition from a milk-based to a fiber rich solid food diet. PedsCom mice developed less peripheral regulatory T cells and Immunoglobulin A, hallmarks of microbiota-driven immune system development. Consistent with defects in maturation, adult PedsCom mice retain high susceptibility to salmonella infection characteristic of young mice and humans. Altogether, our work illustrates how the post-weaning transition in intestinal microbiome composition contributes to normal immune maturation and protection from enteric infection. Accurate modelling of the pre-weaning microbiome provides a window into the microbial requirements of healthy immune development and suggests an opportunity to design microbial interventions at weaning to improve immune system development in human infants.

**One Sentence Summary:** Arresting microbiome development stunts immune ontogeny

## Introduction

Bacteria colonize the intestinal tract at birth, support nutrient metabolism and shape immune system development and function^1^. The early-life microbiome is compositionally distinct from that of older individuals^2^, and these differences have been implicated in increased susceptibility to enteric pathogens in infants^3^. This increased susceptibility to enteric infections has long been thought to be due to immune system immaturity^4^, but has been more recently linked to the immature state of the pediatric microbiome^3^. In humans, the microbiome rapidly shifts its composition, diversity and function from birth to three years of age, after which the microbiome stabilizes and resembles that of an adult microbiome^2^. In mice the developmental progression is more compressed with young mice initiating solid food around day 12-14, fully weaned by day 21 and developing a microbiome which resembles an adult microbiome by day 28^5^. The compositional and functional differences of the early-life microbiome are particularly evident in the pre-weaning stage of life when the infant is solely nourished by maternal milk^6,7^.

Perturbations in the composition of the pediatric microbiome around weaning can alter risk for developing autoimmunity^8^, allergies^9^ and obesity^10^ later in life. Remarkably, the consequences of early-life perturbations persist even if the microbiome is restored to a state resembling healthy subjects^8,9,11^, suggesting that there is a window of opportunity that is critical for host-microbe interactions during ontogeny^12^. Notably, a specific population of regulatory T cells (Tregs), which develop in the gut around weaning, regulate long term immune system health^11^. This is coincident with a spike in pro-inflammatory cytokines tumor necrosis factor alpha (TNF-α) and interferon gamma (IFNγ) expression in the gut, referred to as the ‘weaning reaction’. This process requires the presence of microbes that expand at weaning, and preventing the weaning reaction during ontogeny leads to long-term defects in regulation of inflammatory responses and gut barrier function^11^. Further mechanistic studies are required to understand and leverage the pre-weaning microbiome to support immune system development and health later in life.

Gnotobiotic models, which enable the commensal microbiome to be rationally designed and manipulated, are valuable tools for the study of host-microbiota interactions. For example, studies of gnotobiotic mice demonstrate that specific commensal microbes modulate immune function by inducing the secretion of antimicrobial peptides^13^, development of innate lymphoid cells^14^, peripheral regulatory T cells (pTregs)^15,16^ and IgA production^17^. Gnotobiotic mice designed to model adult microbiomes [e.g., Altered Schaedler’s Flora^18^ (ASF) and Oligo-Mouse-Microbiota-12^19^ (Oligo-MM-12)] reduce the highly complex and diverse adult microbial communities from hundreds of members to a small, defined consortium which facilitates the study of dynamic and complex interactions in fine detail. Such simplified communities can also serve as a platform for the introduction of additional microbes to identify novel functions such as specific systemic antibody targeting of commensals during homeostasis^20^, the ability of pathogenic microbes to alter immune responses to established commensals^21,22^, and the ability of specific commensals to prevent disease^19,23^. However, to date, no published consortium has been designed to specifically model the distinct function of the pre-weaning microbiome.

To experimentally probe the impact of the pre-weaning microbiome on immune function and infection, we developed a tractable gnotobiotic model that mirrors the early-life resident microbiota. Using extensive culture conditions in parallel with metagenomic sequencing, we rationally designed a nine-member microbial consortium which we call Pediatric Community (PedsCom) that represents >90% of the microbial reads present in the intestines of pre-weaning mice. PedsCom is vertically transmitted from dam to pup, and unexpectedly, it remains largely unchanged, retaining the characteristics of a pre-weaning microbiome into adulthood. We found that the consequences of restricting the intestinal microbiome to a pre-weaning state led to a stunting of immune system development and increased susceptibility to neonatal-associated infection.

## Results

### Features of pre-weaning (early-life) microbiomes and construction of a defined consortium functionally similar to infant microbiomes

We first sought to evaluate the development of the intestinal microbiome in our specific pathogen free (SPF) NOD mouse colony from birth to adulthood to determine a pre-weaning timepoint to build a defined consortium. We tracked fecal microbes at days 7, 14, 18, 28, and 42 post birth and assessed microbial composition and diversity by 16S rRNA gene amplicon sequencing of the V4 region. After quality control filtering, the library consisted of 4,241,547 reads clustered into 509 amplicon sequence variants (ASVs) by the DADA2 pipeline^24^. The fecal microbiomes from 7- and 14-day old pre-weaning mice that are exclusively breast-fed were highly similar and dominated by *Lactobacillus*. During the transition from milk to a solid food diet on days 14-18, there was a dramatic shift in the fecal microbiome composition, which by day 18 resembles an adult microbiome (day 42) (**Fig. 1A**). Thus, fecal microbiota derived from 14-day-old mice retain the characteristics of pre-weaning microbiomes yet are poised to undergo rapid maturation during weaning.

**Figure 1.**
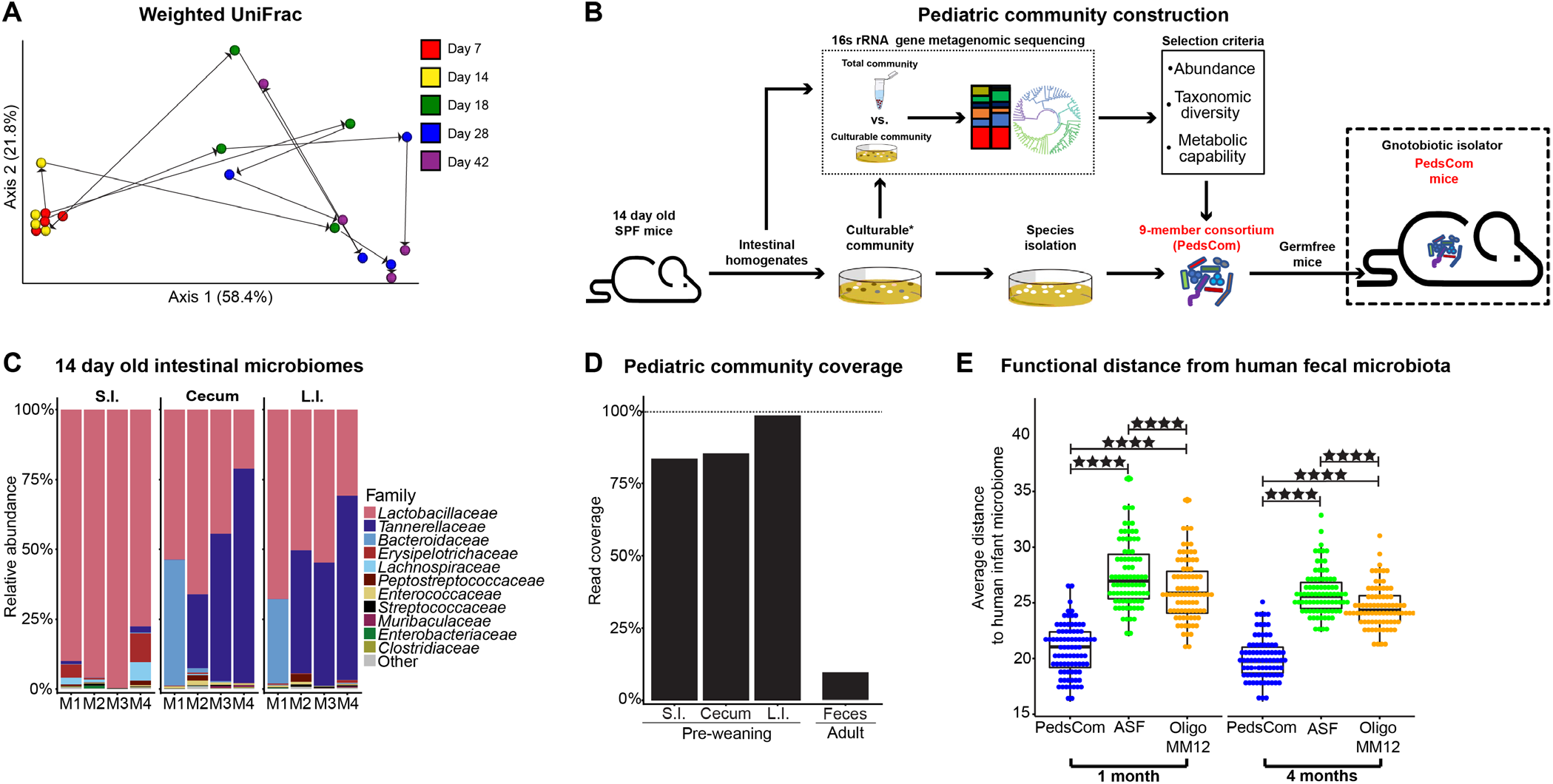
Features of pre-weaning (early-life) microbiomes and construction of a defined consortium functionally similar to infant microbiomes. **A**. Weighted UniFrac Principal Coordinates Analysis (PCoA) of fecal microbiota from SPF NOD mice beta-diversity during ontogeny. Microbial community determined by 16S rRNA gene sequencing. Representative litter shown (N = 4). **B**. Schematic of pediatric community construction. **C**. Intestinal microbiomes of 14-day old donor mice. Family level relative abundance determined by 16S rRNA gene sequencing. N = 4, Data from two litters shown. **D**. Read coverage of PedsCom isolates compared to total organ site (N=4 samples per site). Relative abundance of PedsCom members in the fecal microbiomes of 10-week-old adult SPF NOD mice (N = 19) for comparison. **E**. Euclidean distance comparison of human infant fecal microbiome (N = 84) determined by shotgun metagenomics to the predicated functions of PedsCom, Altered Schaedler Flora (ASF) and Oligo-MM12. Mann-Whitney-Wilcoxon test ****p<0.0001. SPF = Specific pathogen free, S.I. = small intestine, L.I. = large intestine.

We extensively cultured intestinal microbiota from 14-day-old SPF NOD mice, generating a culture collection of microbes from the pre-weaning period (**Fig. 1B**). Tissue from the small intestine, cecum and large intestine from four mice from two litters were collected and homogenized under anaerobic conditions. To maximize yield, these intestinal homogenates were then incubated on diverse media and culture conditions: trypticase soy agar, yeast casitone fatty acid agar, chocolate agar, and environmental conditions: ambient air, ambient air+ 5% CO2, and anaerobic (5% H_2_, 5% CO_2_, 90% N_2_). Individual colonies were isolated and then identified to the species level by 16S rRNA whole gene Sanger sequencing. In parallel to the culture approach, the bacterial composition of the pre-weaning intestinal microbiome was determined by 16S rRNA gene amplicon sequencing of small intestinal, cecal and large intestinal homogenates. After processing, the total library contained 1,311,028 reads from 12 intestinal samples and 67 cultured samples. The small intestine was dominated by *Lactobacillus* spp., which transitioned to a *Bacteroidetes*-dominant (*Tannerellaceae* - *Parabacteroides, Bacteroidaceae* - *Bacteroides*) microbiome in the cecum and large intestine (**Fig. 1C**). There was no significant difference in alpha diversity between intestinal sites (**Fig. S1A**). This pattern differs from what is observed in adult mice, in which diversity increases along the gastrointestinal (G.I.) tract^25,26^. We cultured microbes that account for >99% of the total sequenced reads and 82% of the amplicon sequence variants (ASVs; **Fig. S1B**). Overall, the top 20 ASVs present at each site represented >97% of the read abundance. This low bacterial diversity is a hallmark of the early-life intestinal microbiomes of mice and humans^6^. Altogether, the high percentages of easy-to-cultivate bacterial species and the low diversity of pre-weaning microbiomes provided an opportunity to construct a defined consortium to model the pre-weaning microbiome.

We hypothesized that a defined consortium composed of microbes isolated from pre-weaning mice would effectively colonize the gastrointestinal tract of germfree C57BL/6 mice, be vertically transmitted from dam to offspring, and recapitulate the functions of early-life intestinal microbiomes in progeny. A consortium of nine isolates designated as Pediatric Community (PedsCom) (**Table 1**) was selected based on relative abundance across the intestinal sites, taxonomic diversity and predicted metabolic capacity. These nine bacteria represent a remarkably high proportion of the microbial sequences and predicted functions present in the small intestine, cecum, and large intestine (83%, 85%, and 98%, respectively) but only 9.6% of the relative abundance of conventionally raised adult fecal microbiomes (**Fig. 1D and S1C**).

**Tbl.1 –.**
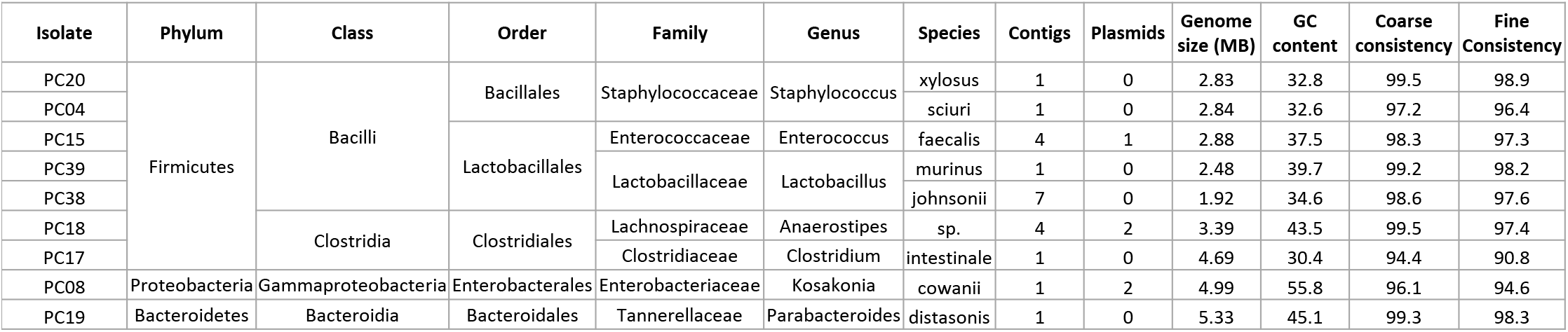
Summary of PedsCom consortia

To investigate the functional capabilities of the PedsCom consortium, we performed whole genome sequencing of each species. Isolates were sequenced with the Oxford Nanopore and Illumina Miseq platforms and the combined reads assembled with the Unicycler pipeline^27^. Genomes were annotated using the RAST algorithm through the PATRIC webserver^28,29^. In aggregate, the genes of the PedsCom consortium consisted of a total of 30,292 coding sequences (CDS) that were assigned to 14,988 molecular functions [KEGG Orthology groups; (KOs)] ^30^.

Comparison to two other gnotobiotic models (ASF and Oligo-MM12) revealed taxonomic and likely functional differences. ASF is an eight-member consortium developed by Orcutt and colleagues in 1978, which has been extensively used in gnotobiotic studies^18^. Oligo-MM12 is a more recent genome-guided design of the mouse intestinal microbiome with microbes derived from adult mice^19^. Taxonomically, PedsCom is composed of a larger proportion of isolates from the class *Bacilli (Enterococcus faecalis, Lactobacillus johnsonii, Lactobacillus murinus, Staphylococcus sciuri*, and *Staphylococcus xylosus*) compared to ASF or Oligo-MM12. The increase in *Bacilli* mirrors the increased prevalence of this class in infants compared to adults^6,7,31^. *Clostridium intestinale* is a member of *Clostridiaceae* (clostridium cluster I), a family that is strongly associated with the early-life microbiome that decreases during maturation^32^, and is absent in ASF and Oligo-MM12. In PedsCom, the *Lachnospiraceae* lineage of *Clostridia* (cluster XIVa) is also represented by *Anaerostipes* sp. PC18; potentially a novel species closely related to *Anaerostipes caccae* (average nucleotide identity - 92.16%). The *Bacteroidetes* lineage is represented by *Parabacteroides distasonis*, comparable to ASF, which contains the related *Parabacteroides goldsteinii*. Lastly, PedsCom contains an *Enterobacteriaceae* isolate, *Kosakonia cowanii*, a family of microbes absent in both ASF and Oligo-MM12. *Enterobacteriaceae* is a critical component of modeling the pre-weaning microbiome with levels reaching ~20% relative abundance in human neonates^33^.

Based on the taxonomic associations of PedsCom members with the early-life microbiome, we hypothesized that PedsCom is more functionally similar to human early-life microbiomes than either ASF or oligo-MM12. We compared the three consortia by analyzing annotated Kyoto Encyclopedia of Genes and Genomes (KEGG) orthologs (KOs)^30^. Briefly, open reading frames in each isolate were assigned to a KO using the KEGG automatic annotation server (KAAS)^30^. The KO abundances of the three consortia were compared to a published metagenomic data set of fecal microbiomes from 84 human infants at one and four months of age^34^. The PedsCom community was more similar to the human infant fecal microbiota at both one and four months old by average Euclidean distance than either ASF or OligoMM-12 (**Fig. 1E**), indicating that the functional repertoire of PedsCom more closely resembles that of pre-weaning age microbiomes of human infants.

### Community dynamics of PedsCom consortium is restricted during weaning

We next investigated whether the nine-member PedsCom consortium stably colonizes germfree mice and the effect of weaning on community composition. To generate gnotobiotic PedsCom mice, 1×10^9^ colony-forming units (CFUs) of each PedsCom isolate were orally gavaged into 6-week-old female C57BL/6 germ-free (GF) mice (n=3). All nine microbes were recovered from the fecal microbiome one day post-gavage. From day 2 to day 28, the fecal microbiomes were dominated by *P. distasonis* and *K. cowanii*. The species *L. johnsonii, S. xylosus, and S. sciuri* were below 0.1% relative abundance one-day post inoculation, and undetectable in the fecal microbiome by day seven. In contrast, *C. intestinale*, which was present at low abundance on day 1 (0.029%), increased 10-fold by day 7 (0.31%) and 60-fold to 1.74% by day 21. Overall, 7 of 9 members PedsCom consortium of microbes were detected in the initial survey of the fecal microbiomes of germfree mice gavaged as adults, and these microbes formed a stable community that resembles a normal pre-weaning microbiome (**Fig. 2A**). The two microbes that appeared to be lost (*S. xylosus, and S. sciuri*) were detected in progeny born to these adult PedsCom founders indicating successful colonization and vertical transmission of all 9 PedsCom microbes.

**Figure 2.**
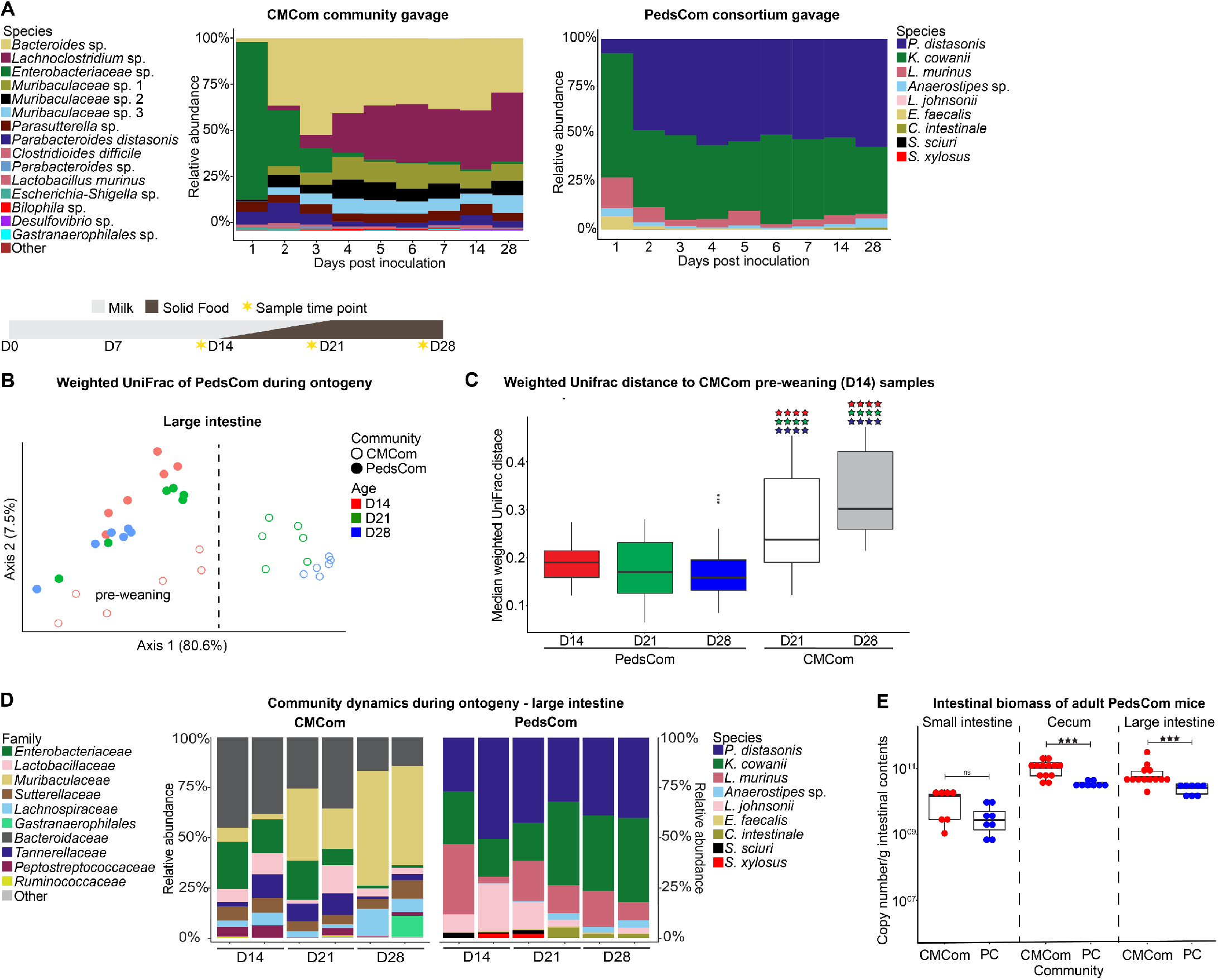
Community dynamics of PedsCom consortium is restricted during weaning. **A**. Representative species level fecal microbiota of ex-germfree female C57BL/6J mice gavaged with PedsCom consortium (N = 6 mice, 2 independent experiments) or CMCom community (N = 3 mice). **B**. Comparison of cecal and large intestinal microbiota of PedsCom and CMCom mice at 14, 21 and 28 days old by weighted UniFrac PCoA beta-diversity of 16S rRNA gene sequencing. Dashed line indicates separation of CMCom pre-weaning and post-weaning samples. N = 6 samples per timepoint. **C**. Median weighted UniFrac distance comparison of day 14 CMCom large intestine microbiota (pre-weaning) to all PedsCom and post-weaning (D21, D28) CMCom microbiota. Color of stars indicates PedsCom timepoint compared (D14 = red, D21 = green, D28 = blue). N = 6 samples per timepoint. Mann-Whitney-Wilcoxon test ****p<0.0001. **D**. Relative abundance of PedsCom and CMCom mice microbiota during ontogeny in the large intestine. Two representative litters for each community and timepoint. **E**. Intestinal biomass of adult CMCom and PedsCom mice (16S rRNA gene copies per gram intestinal contents). (PedsCom – N = 8 per tissue, CMCom - S.I. N = 7, Cecum and L.I, N = 13). Mann-Whitney-Wilcoxon test **p<0.01, ***p<0.001.

To generate a control line of mice harboring an undefined, complex community, the cecal contents of a conventionally housed 6-week-old adult mouse was gavaged into female GF mice (n=3) and maintained in a separate gnotobiotic isolator. We designated this closed but complex adult community as the Complex Mature Community, CMCom. The fecal microbial communities from CMCom gavaged mice exhibited substantial changes in relative abundance during the first week post-colonization (**Fig. 2A**). *Enterobacteriaceae* initially dominated the fecal microbial community with >80% relative abundance by day 1, but by the end of the first week, microbes from the *Bacteroidetes* and *Firmicutes* became the dominant taxa. This pattern recapitulated normal development of the microbiome although at a much-accelerated time scale.

The stability of PedsCom during adult colonization led us to investigate whether the PedsCom community would undergo weaning-induced changes in relative abundance in the progeny of PedsCom mice. Age-determined progression was observed in CMCom mouse intestinal samples, demonstrating a clear developmental arc of community composition from days 14 to 28. Unexpectedly, we found no clear separation of intestinal samples by age in PedsCom mice by weighted UniFrac analysis (**Fig. 2B, Fig. S2A,C**). In fact, PedsCom samples from all age groups clustered more closely with 14-day old microbiomes of CMCom mice, suggesting persistent similarity to the pre-weaning microbiome (**Fig. 2C**). Further supporting this observation, the relative abundances of dominant PedsCom members remained remarkably stable in the small intestine, cecum and large intestine during the transition from a milk-based diet to a solid food diet [*P. distasonis* (36.04% to 34.46%), *K. cowanii* (22.03% to 32.45%) and *L. murinus* (13.96% to 18.92%] (**Fig. 2D and S2B,D**). Modest shifts in the relative abundances of taxa in the large intestines included increases in *Anaerostipes* (0.49% to 1.57%), *C. intestinale* (0.01% to 2.73%) and reduction of *L. johnsonii* (1.97% to 0.04%), *S. xylosus* and *S. sciuri* (0.02% to undetectable) by week four. CMCom and PedsCom had several taxa in common (e.g., *P. distasonis, L. murinus, E. faecalis*) in the cecum and large intestine, especially at earlier time points. However, in CMCom these taxa decreased in abundance and were replaced by *Muribaculaceae*, *Bacteroidaceae* and *Lachnospiraceae* during the transition to solid food from 14 to 28 days old. In contrast, relative abundances of these bacteria in PedsCom mice remained unchanged into adulthood (**Fig. S2E**). We also observed significantly lower biomass in the cecum (3.4 × 10^10^ vs. 1.0 × 10^11^) and large intestine (2.6 × 10^10^ vs. 5.6 × 10^10^) of PedsCom mice compared to CMCom, which is consistent with lower biomass in pre-weaning compared to adult cecal microbiota^35^ (**Fig. 2E**). Taken together, the PedsCom nine-member consortium of microbes derived from the pre-weaning microbiota is vertically transmitted from dam to pups and remains stable through the weaning period into adulthood. This ‘locking in’ of the PedsCom community, discovered serendipitously, provides an opportunity to investigate the immunologic and physiologic impacts of progression from a pre-weaning to adult microbiome.

### Restriction of intestinal microbiome maturation stunts cellular and humoral immune system development

We used the PedsCom model to test whether components of the immune system that depend upon normal microbiome maturation are impacted by locking in a pre-weaning intestinal community. We hypothesized that arresting microbiome maturation at weaning would stunt the development of those components of the immune system that develop during and soon after weaning. We focused on CD4^+^Foxp3^+^Rorγ^+^ peripheral Tregs (pTregs) which are induced by specific commensal microbes and food antigens during weaning and are required to maintain intestinal homeostasis^15,16,36^ (**Fig. 3A**). As expected, pTreg proportions increased in CMCom mice from 14-28 days old, with a sharp rise occurring post-weaning^15,16^ (**Fig. 3B**). In contrast, pTregs number and frequency was considerably lower but not absent in PedsCom mice. The partial induction of pTregs in PedsCom mice suggests that radical shifts of abundance or novel introductions of microbes colonizing the gut is not required to induce this cell population at weaning. However, the pTreg “ceiling” was considerably lower in PedsCom mice post-weaning, such that as the overall proportion of pTregs continues to increase in PedsCom mice into adulthood, the populations remained significantly lower than that of CMCom (**Fig. 3C**).

**Figure 3.**
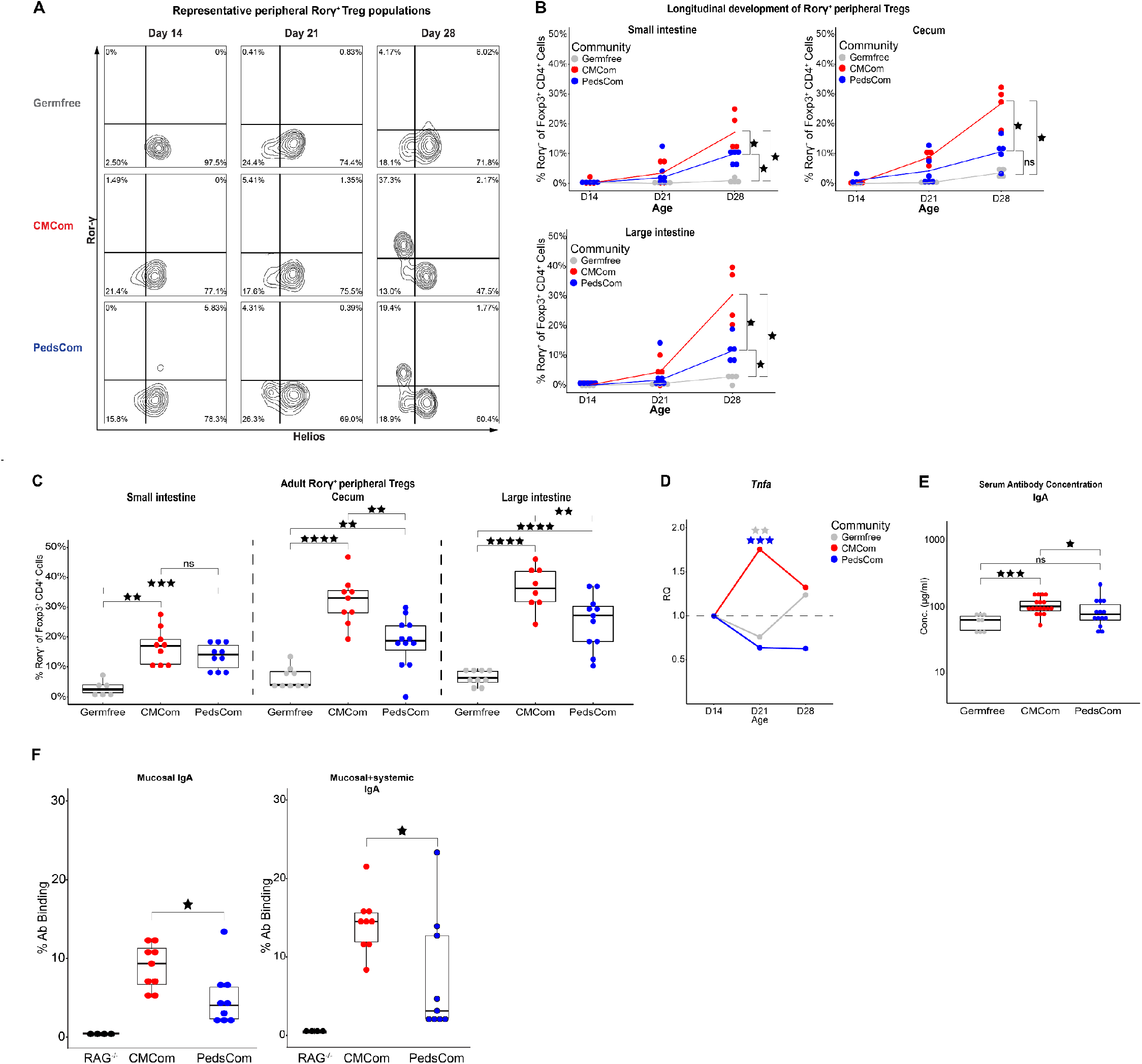
Restriction of intestinal microbiome maturation stunts cellular and humoral immune system development. **A**. Representative flow cytometry plot of CD4^+^Foxp3^+^Rory^+^Helios^-^ peripheral Treg (pTreg) cell population. Cells were gated on CD45^+^TCRβ^+^CD19^-^CD8^-^CD4^+^Foxp3^+^. **B**. Percentage of pTreg cells in the small intestine, cecum and large intestine lamina propria of germfree, PedsCom and CMCom mice during ontogeny. N = 3 for day 21 germfree cecum, N ≥ 4 per remaining community and timepoint. Mann-Whitney-Wilcoxon test *p<0.05, **p<0.01, ***p<0.001. **C**. Percentage of pTreg cells in the small intestine, cecum and large intestine lamina propria of germfree, PedsCom and CMCom 7–12-week-old adult mice. N = 3 for germfree S.I., N ≥ 8 per remaining tissue and timepoint. **D**. Tumor necrosis factor alpha (*TNFα*) transcript levels in distal small intestinal tissue of germfree, PedsCom and CMCom mice during weaning. Relative quantitation (RQ) represents the fold change at day 21 and day 28 relative to mean baseline transcript levels at day 14 in each community. Day 21 represents pooled data of mice sacrificed at day 21 and day 22 post birth. Germfree (N: D14 = 3, D21 = 7, D28 = 8), PedsCom (N: D14 = 4, D21 = 7, D28 = 4), CMCom (N: D14 = 7, D21 = 13, D28 = 5). Independent sample t-test **p<0.01, ***p<0.001. **E**. Serum IgA concentration in germfree, PedsCom and CMCom 6–12-week-old adult mice. (N: Germfree = 8, CMCom = 18, PedsCom = 14). **F**. Percent mucosal and systemic IgA binding to PedsCom and CMCom 7-10-week-old adult fecal microbes. (N: RAG^-/-^ = 4, CMCom, PedsCom = 9). Mann-Whitney-Wilcoxon test *p<0.05, **p<0.01, ***p<0.001.

The development of pTregs in gut tissues coincides with a spike in pro-inflammatory cytokines in the gut, known as the ‘weaning reaction’. Consistent with conventionally housed mice^11^, *Tnfa* gene expression was elevated at day 21 in CMCom but not PedsCom small intestine relative to day 14 levels (**Fig. 3D**). This data indicates that the PedsCom microbes do not induce the characteristic *Tnfa* inflammatory response at weaning. We also analyzed other lymphoid cell populations in the lamina propria of the small intestine, cecum, and large intestine. Overall number and percentage of αβ and γδ T cells and B cells were similar between PedsCom and CMCom mice. Further, we found similar numbers and percentages of CD4^+^ and CD8^+^ T cells.

Germfree mice have lower levels of serum IgA, and higher levels of serum IgE due to loss of commensal stimulation^37^. These effects become evident soon after weaning when maternal IgA wains and nascent endogenous IgA becomes the dominant mucosal antibody^38^. To investigate the effect of PedsCom on serum immunoglobulin (Ig) levels, quantitative ELISA was performed on adult PedsCom mice with germfree and CMCom as control groups. Serum IgA concentration (78.0 μg/ml) were significantly lower in PedsCom mice compared to CMCom mice (101.7 μg/ml) and similar to germfree mice (63.8 μg/ml) (**Fig. 3E**). Notably, IgE concentration was significantly higher in PedsCom mice compared to CMCom controls (30.0 ng/ml vs. 13.7 ng/ml) (**Fig. S3A**). Diminished serum IgA and elevated serum IgE are both consistent with decreased microbial stimulation of the immune system during development.

Since PedsCom colonization leads to reduced systemic IgA, we hypothesized that PedsCom microbes may also induce less intestinal IgA capable of binding to these commensal microbes. Indeed, microbial flow cytometry (mFLOW) analysis of fecal microbiota (**Fig. S3B**) showed that IgA binds to a lower percentage of the fecal microbes in PedsCom compared to CMCom mice (4.1% vs 9.3%) (**Fig. 3F**). Further incubating fecal microbiota with autologous serum antibodies showed less binding of IgA to the PedsCom microbes compared to CMCom (**Fig. 3F**). Notably, serum IgA did not appreciably increase percent binding to PedsCom fecal microbes compared to feces alone (3.1%), though a slight increase was seen in CMCom samples (14.5%). In summary, lower serum IgA levels in PedsCom mice correspond to less mucosal IgA targeting of the PedsCom consortia members. Reduced intestinal pTreg populations and commensal-mediated antibody levels indicates that the restriction of PedsCom mice intestinal microbiota to a pre-weaning configuration stunts intestinal and systemic immune development.

### PedsCom mice are highly susceptible to *Salmonella* infection

Since PedsCom mice have a ‘locked in’ pre-weaning microbiome and defects in immune system development, we next assessed the ability of PedsCom mice to respond to salmonella infection, a leading cause of pediatric mortality^39^, whose susceptibility is linked to features of the pediatric intestinal microbiome ^3,40^. To determine if adult mice harboring PedsCom maintain the high susceptibility of neonatal mice to enteric pathogens, we orally challenged PedsCom, germfree and CMCom adult mice with 5×10^8^ CFUs of wild-type *S. typhimurium* SL1344. Germfree mice were highly susceptible to salmonella infection with 100% mortality occurring within 3 days (**Fig. 4A**). CMCom mice were resistant with 100% survival 7 days post infection (d.p.i.). PedsCom mice however had an intermediate survival. The germfree cohort had severe diarrhea 1 d.p.i., likely reflecting overwhelming intestinal inflammation^41^, which was not evident in PedsCom mice or CMCom mice, consistent with histopathology which revealed a near-complete destruction of the intestinal epithelial layer only in the germfree mice. The increased mortality of PedsCom-colonized adult mice compared to CMCom suggests that the PedsCom mice lack protective mechanisms found in mice colonized with adult intestinal communities.

**Figure 4.**
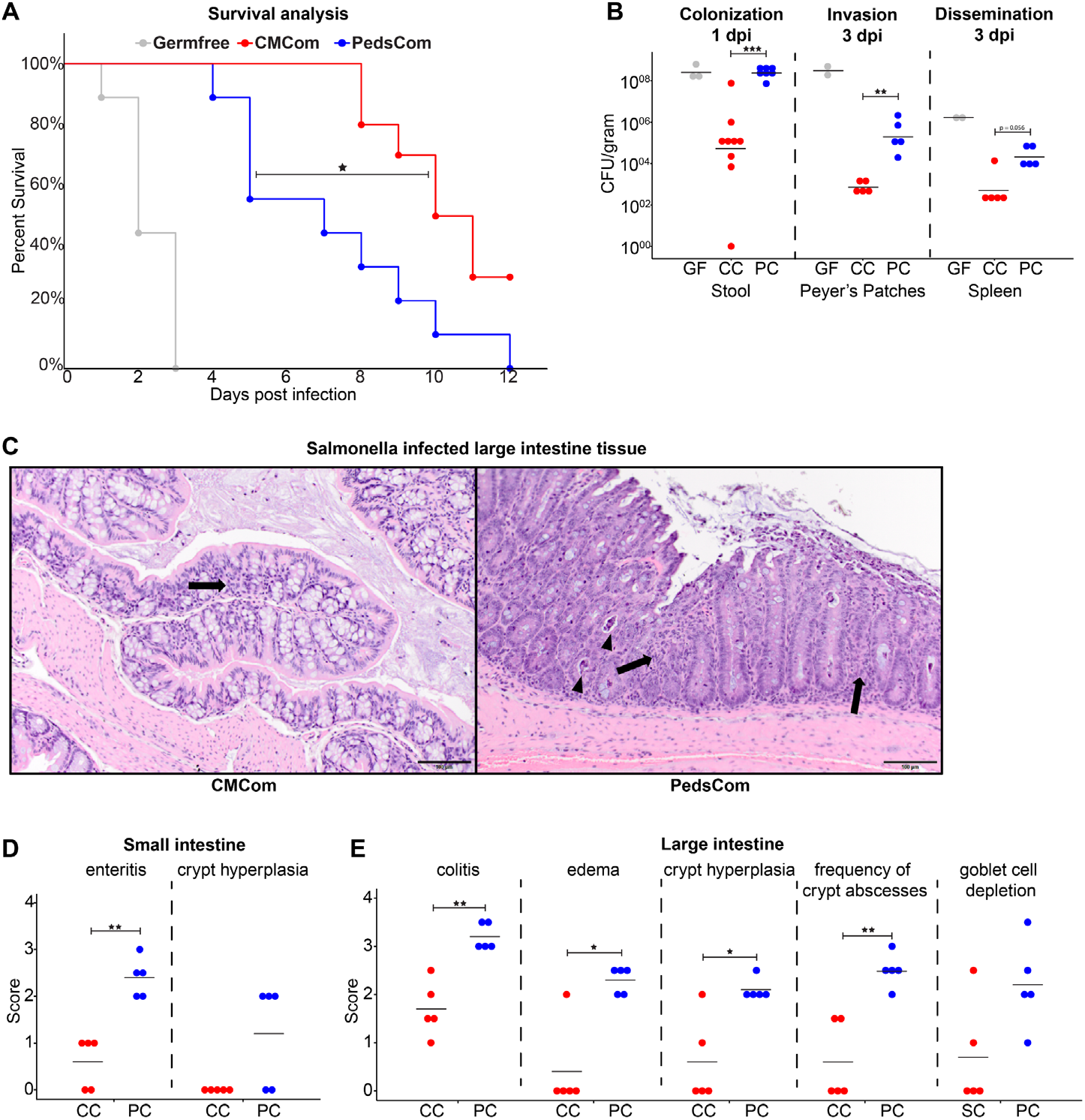
Increased susceptibility of PedsCom gnotobiotic mice to *Salmonella typhimurium* infection. **A**. Percent survival of adult PedsCom, CMCom and GF mice infected with *Salmonella typhimurium* SL1344. Adult mice were 10-17 weeks of age. Germfree (N = 9), PedsCom (N = 9), and CMCom (N = 10). Log-rank Mantel-Cox test *p<0.05. Data from two independent experiments. **B**. Salmonella burden 1 day post infection in feces [Germfree (N = 3), PedsCom (N = 9), and CMCom (N = 7)], Peyer’s patches and spleen 3 days post infection [Germfree (N = 2), PedsCom and CMCom (N = 5)]. CFU per gram feces or indicated tissue. GF = Germfree, CC = CMCom, PC = PedsCom. **C**. Pathologic features of CMCom and PedsCom large intestine tissue 3 days post salmonella infection. Arrows indicate minor inflammatory infiltrates within the mucosa and arrowheads indicate crypt abscesses. (H&E staining, 200x magnification, scale bar: 100 μm). **D and E**. Pathological scores of small and large intestine tissue 3 days post salmonella infection in PedsCom and CMCom mice (N = 5 per tissue). Mann-Whitney-Wilcoxon test *p<0.05, **p<0.01, ***p<0.001.

To investigate the pathogenesis of salmonella infection in PedsCom mice, we compared *S. typhimurium* burden in the gut, invasion into Peyer’s patches and systemic dissemination between the three groups. We found a ~3 log increase in CFUs per gram of feces at day one in PedsCom mice compared to CMCom controls (**Fig. 4B**) indicating increased *S. typhimurium* burden of the distal intestines in PedsCom mice. The increased salmonella burden of PedsCom mice relative to CMCom is consistent with the phenotype reported in young SPF mice and studies of transplanted complex neonatal and pre-weaning microbiota into adult germfree mice^3^. In contrast, *S. typhimurium* burden in PedsCom mice was comparable to germfree controls, despite a large discrepancy in survival between these two groups, indicating that high salmonella levels in the distal intestines alone do not account for differences in survival. PedsCom mice had significantly higher *S. typhimurium* CFUs in the Peyer’s patches and trended higher in the spleen of PedsCom mice compared to CMCom mice (**Fig. 4B**). Thus, PedsCom mice are more susceptible to pathogen invasion of the Peyer’s patches and subsequent systemic invasion than CMCom mice, likely explaining the increased mortality. However, compared to germfree controls, PedsCom partially prevents invasion of Peyer’s patches and systemic spread, despite similar *S. typhimurium* fecal densities.

To assess the level of intestinal pathology caused by salmonella infection in PedsCom and CMCom, we investigated small intestine and large intestine pathology. Comparison of histological sections of uninfected PedsCom and CMCom mice found no obvious differences in intestinal development between the colonized groups (**Fig. S4**). The most common lesions in infected mice included inflammation characterized by inflammatory cells (predominantly neutrophils, lymphocytes, and histiocytes) expanding in the lamina propria of the mucosa and in the most severely affected animals, effacing and replacing crypts (**Fig. 4C**). Secondary changes included attenuation of the mucosa, crypt hyperplasia, edema, crypt abscesses (degenerate leukocytes and necrotic debris within crypt lumens), or necrotic debris in the lumen of the intestine. The large intestine exhibited more severe damage than the small intestine, consistent with salmonella pathogenesis in streptomycin treated mice^42^. Small intestine histology revealed significantly worse enteritis in all PedsCom mice (**Fig. 4D**). Consistent with the small intestine data, we found significantly higher colitis, edema, crypt hyperplasia scores, and an increased frequency of crypt abscesses in infected PedsCom large intestines (**Fig. 4E**). Overall, these data suggests that adult PedsCom mice retain the susceptibility to salmonella infection characteristic of pre-weaning mice, demonstrating the need for intestinal microbiome maturation in preventing enteric infection.

## Discussion

In this study, we developed a microbial community, ‘PedsCom’ that recapitulates the relative abundances (>90%), taxonomic diversity and functional capability of pre-weaning intestinal microbiomes. Using these mice, we made two important findings. First, PedsCom unexpectedly retains the characteristics of a pre-weaning microbiome into adulthood. Second, restriction of the intestinal microbiome to a pre-weaning-configuration leads to stunting of immune system development.

The finding that the most prevalent microbes pre-weaning are unaffected by the dietary change to solid food provokes the question of whether introduction of solid food alone is insufficient to drive maturational shifts of the relative abundance of the predominant early-life microbes in the gut. Further, it suggests that exogenous or low-abundant microbes may be required for microbiome maturation to occur at weaning. Interestingly, complex pre-weaning communities of microbes introduced into adult germfree mice also retain their pre-weaning configuration in the presence of solid food^3^, suggesting that the timing of microbial exposure at weaning may be necessary for intestinal microbiome maturation. This would signify that the weaning period is a critical and complex window for the introduction of new microbes and maturation of the microbiome. Further, the weaning period may be an ideal time to successfully introduce probiotics which may better engraft and influence immune system development.

Restriction of the intestinal microbiome to a pre-weaning-configuration leads to stunting of immune system development illustrated by: 1) decreased pTreg expansion in intestinal tissues, 2) loss of the *Tnfa*-mediated weaning reaction, 3) lower serum IgA and 4) lower fecal IgA binding to commensal microbiota, 5) abnormally high serum IgE and 6) increased susceptibility to enteric infection. These findings highlight that weaning provides a developmental window in which exogenous or low-abundance endogenous microbes required for microbiome maturation drive development of cellular and humoral immunity.

A question arises as to what factors are missing in PedsCom that are necessary for robust pTreg generation and the weaning reaction. Several studies have shown a role of microbially-derived short chain fatty acids (SCFAs), derived from catabolism of plant-derived complex carbohydrates found in solid food, in the weaning reaction^11^ and subsequent intestinal Treg induction^43–45^. In particular, colonic Treg expansion has been associated with *Clostridia* of group XIVa and IV^46,47^, microbes associated with SCFA production that increase in abundance post-weaning. Though the PedsCom consortium includes pTreg inducing isolates, it is possible that the lack of clostridial species acquired post-weaning limits full pTreg expansion and subsequent immune maturation. Thus, a possible mechanism explaining the absence of a weaning reaction and partial induction of Tregs is that the members of PedsCom, dominant when milk is plentiful, lack the metabolic capability to efficiently metabolize more complex carbohydrates and therefore produce fewer SCFAs.

The increased susceptibility of adult PedsCom mice to salmonella infection could be due to stunting of immune maturation or, alternatively, to the restriction to a pre-weaning intestinal microbiome. Compared to CMCom controls, *S. typhimurium* causes severe intestinal epithelial damage and immune cell infiltration in PedsCom mice, quickly becoming a systemic infection. It is possible that this phenotype is due to a decrease in barrier function and increased pro-inflammatory responses associated with disruption of the weaning reaction, pTreg generation and normal gut immune maturation^11^. A predisposition to intestinal inflammation can increase salmonella burden, since *S. typhimurium* can gain a competitive advantage in the gut under inflammatory conditions^48^. Notably, commensally-induced pTregs may reduce pro-inflammatory cytokine production during salmonella infection, reducing disease severity and systemic bioburden^49^.

It is also possible that the low complexity of the PedsCom consortium is responsible for the susceptibility to salmonella infection. Though complete early-life microbiomes have lower complexity than adult microbiomes, the increased susceptibility of PedsCom to salmonella infection in mice may be due to the limited number of taxa present in a defined consortium that can compete with the invading pathogen. Compared to SPF controls, ASF colonized adult mice have also been shown to be more susceptible to salmonella infection, with the low complexity of the consortium described as the cause^50^. However, when intact pre-weaning intestinal communities were transplanted into adult germfree mice, they remained susceptible to *S. typhimurium* infection and mortality, while mice colonized with adult communities were resistant^3^. In addition, dietary supplementation that shifted adult communities into an pre-weaning configuration increased *S. typhimurium* burden upon infection^3^, indicating that adult-associated microbes are responsible for reducing salmonella infection and not the absolute number of species. Furthermore, Brugiroux *et. al* ^19^ showed that the addition of 5 supplementary microbes from ASF to Oligo-MM12 (now an 18-member consortium) did not alter *S. typhimurium* burden compared to Oligo-MM12 alone. In contrast, the addition of three facultative anaerobes (*E. coli* Mt1B1, *Streptococcus danieliae* ERD01G and *S. xylosus* 33-ERD13C, referred to as ‘FA3’), now a 15-member consortium, reduced *S. typhimurium* burden in Oligo-MM12 mice. This indicates that features of microbes introduced, as opposed to absolute numbers, is responsible for reducing *S. typhimurium* burden. The ability of certain microbes to decrease infection and not others highlights what one would intuitively expect; that just increasing the number of species present will not necessarily decrease pathogen burden. It is likely that the susceptibility to death from salmonella infections in adult PedsCom mice is due to both stunted immune development leading to a pro-inflammatory environment during infection and the lack of adult-associated taxa to effectively occupy available niches and compete for resources.

Our study has defined a consortium (PedsCom) as a model for investigating the unique function of the pre-weaning intestinal microbiome and its impact on host functions. Critically, PedsCom’s resistance to weaning-associated microbial shifts and stunted immune maturation allows it to serve as a model to elucidate host functions that require transition to an adult-associated microbiome. The PedsCom consortium provides a novel tool for the assessment of features of immunomodulatory microbes required during the weaning to prevent the pathologies associated with microbiome perturbations during this period in human development. Accurate modelling of the pre-weaning microbiome provides a high-impact opportunity to design microbial interventions at weaning to improve immune system development and long-term immune health.

## Acknowledgements

The authors thank Kyle Bittinger, Ceylan Tanes, William Bailis, Matthew Weitzman, Laurence Eisenlohr and Joseph Zackular for helpful discussions. The authors also thank Dmitri Kobuley and Michelle Albright of the PennCHOP microbiome program gnotobiotic mouse facility at Hill Pavilion.

## Author Contributions

Conceptualization, J.L., P.J.P. and M.A.S.; Methodology, J.L., I.E.B, P.J.P. and M.A.S.; Investigation, J.L., J.G., S.M., L.D., T.D., M.L. and M.W.D.; Writing – Original Draft, J.L., P.J.P. and M.A.S.; Writing – Review & Editing, J.L., J.G., S.M., M.L., P.J.P., and M.A.S.; Funding Acquisition, P.J.P. and M.A.S.; Resources, I.E.B, P.J.P. and M.A.S.; Supervision, P.J.P. and M.A.S.

---

**Supplemental Figure 1.**
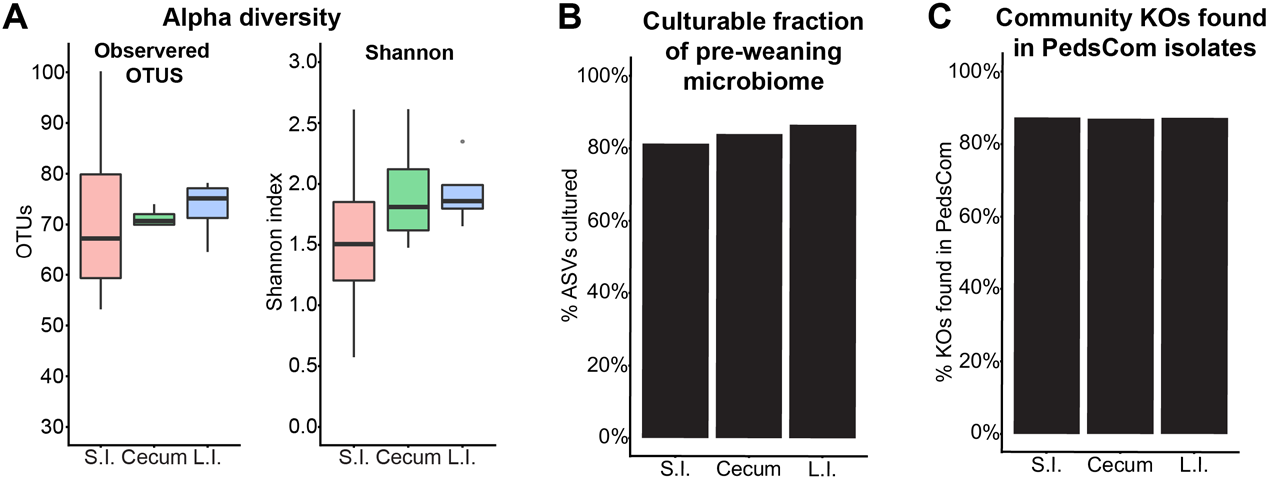
Low complexity of pre-weaning (early-life) microbial community enables recapitulation of microbial abundance and function in limited consortium. **A**. Alpha diversity of the intestinal microbiomes from 14-day-old donor mice (N = 4 samples per tissue site). **B**. Percentage of the total amplicon sequence variants (ASVs) present in each tissue that were recovered from bacterial culture plates. **C**. Percent of KOs represented by PedsCom consortium in complete sequenced community in small intestine, cecum, and large intestine of day 14 donor mice.

**Supplemental Figure 2.**
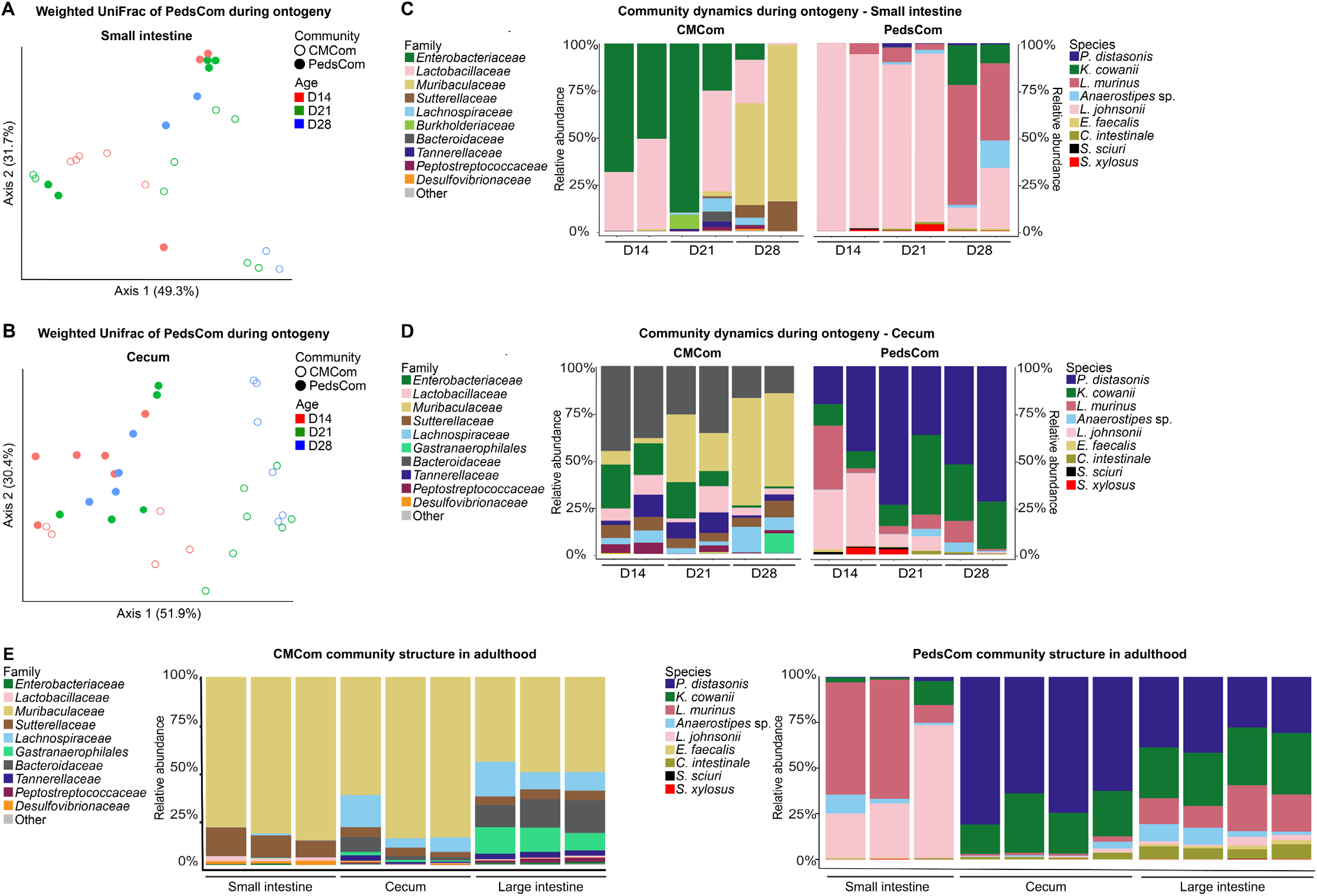
Community dynamics of PedsCom consortium is restricted during weaning. **A**. Weighted UniFrac PCoA beta-diversity of small intestinal microbiota of PedsCom (N: D14 = 3, D21 = 5, D28 = 2) and CMCom (N: D14 = 5, D21 = 8, D28 = 2) mice during ontogeny. **B**. Species and family level 16S rRNA relative abundance of PedsCom and CMCom mice microbiota during ontogeny in the small intestine. Two representative litters for CMCom per timepoint, one litter represented in PedsCom. **C**. Weighted UniFrac PCoA beta-diversity of cecal microbiota of PedsCom (N: D14 = 6, D21 = 5, D28 = 4) and CMCom (N: D14 = 5, D21 = 7, D28 = 6) mice during ontogeny. **D**. Species and family level 16S rRNA relative abundance of PedsCom and CMCom mice microbiota during ontogeny in the cecum. Two representative litters for communities per timepoint. **E**. Representative litter of species and family level 16S rRNA gene relative abundance of adult PedsCom and CMCom intestinal microbiota.

**Supplemental Figure 3.**
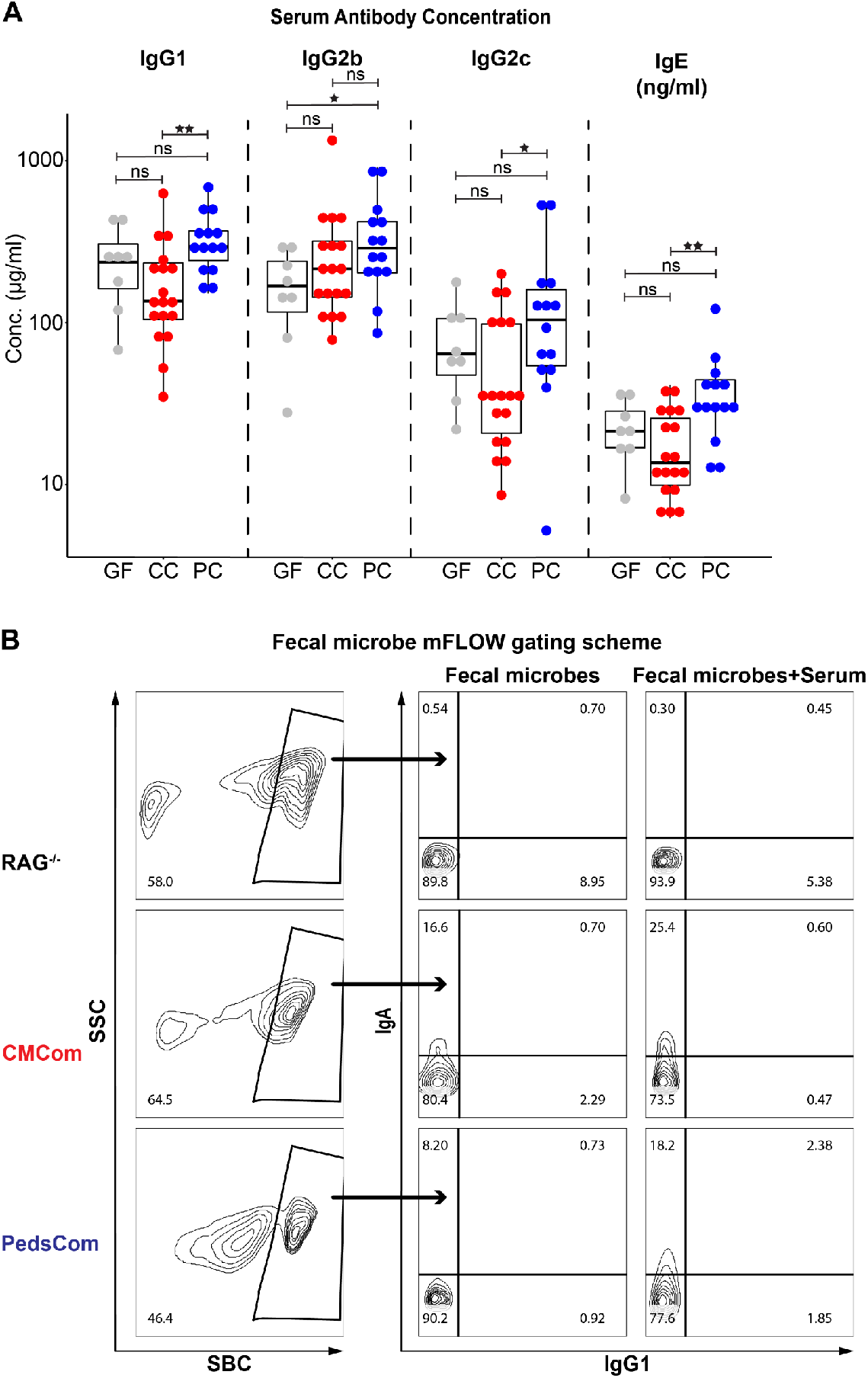
PedsCom consortium leads to increased systemic immunoglobulin levels. **A**. ELISA quantification of serum IgG1, IgG2b, IgG2c and IgE concentration in 7-12-week-old adult PedsCom and CMCom mice. N values: (Germfree = 8, CMCom = 18, PedsCom = 14). GF = Germfree, CC = CMCom, PC = PedsCom. **B**. Representative gating scheme of microbial flow cytometry (mFLOW) analysis of immunoglobulin binding to fecal microbes.

**Supplemental Figure 4.**
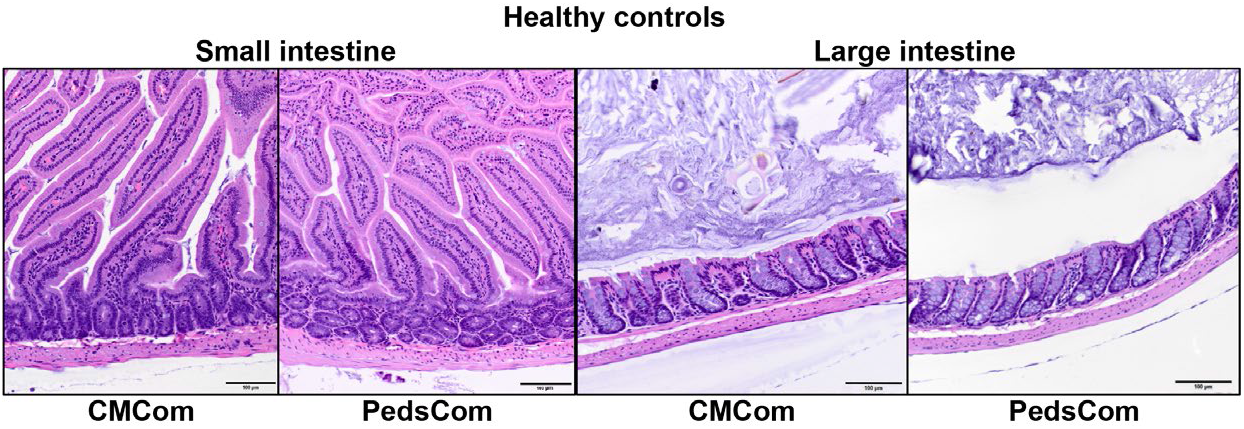
PedsCom mice develop similar intestinal morphology to CMCom mice. Comparison of H&E stained healthy, adult CMCom and PedsCom small and large intestine tissue sections. Representative samples shown, N = 3 (200x magnification, scale bar: 100 μm).

## Methods

### Mice

Specific pathogen free NOD mice were bred and housed in ARC animal facility at Children’s Hospital of Philadelphia. ARC housed mice were fed LabDiet 5058 (Cat# 0007689, MO, USA) ad libitum. Germfree and gnotobiotic C57BL/6J mice were derived and housed in the Hill Pavilion gnotobiotic mouse facility, University of Pennsylvania. Germfree and gnotobiotic mice are housed in flexible film isolators (Class Biologically Clean (CBClean), WI, USA). Hill housed mice were fed autoclaved LabDiet 5021 (Cat# 0006540) ad libitum. Germfree and gnotobiotic mice were caged on autoclaved Beta-chip hardwood bedding (Nepco, NY, USA). Sterility checks were regularly performed on isolators twice a month and additionally prior to any transfer of animals. Freshly collected pellets were cultured on brain heart infusion (BHI) (Oxoid, UK), NB1, and Sabouraud media for 65-70 hours at 37°C under aerobic and anaerobic conditions with positive and negative control samples. Isolator sterility was confirmed externally every 3-4 months by Charles River Laboratories (NJ, USA).

### Bacterial strain isolation and gavage conditions

Whole small intestine, cecum, and large intestine tissues were collected from 14-day old mice and homogenized under anaerobic conditions (90% N_2_, 5% CO_2_, 5% H_2_) in sterile, reduced 0.1% L-cysteine, phosphate buffered saline (rPBS). A sample of the organ homogenates was collected and stored at −80°C for sequencing. Serial dilutions of homogenates were plated on four bacterial growth media: Tryptic Soy Agar (Becton-Dickinson (BD), MD, USA), Chocolate Agar (BD), Laked Blood Agar with 100 mg/L Kanamycin, 7.5 mg/L Vancomycin (BD), and Yeast Casitone Fatty Acid (YCFA) prepared as previously described^51^. Plates were cultured under three different growth conditions: anerobic at 37°C for five days, ambient air at 37°C for three days, ambient air supplemented with 5% CO_2_ at 37°C for three days. Colony isolates were restreaked for purity and preliminary identification performed by 16S rRNA Sanger sequencing. Isolates were stored at −80°C in 25% glycerol, BHI media. Whole plate scrapings were collected for sequencing. PedsCom bacteria were grown under the following conditions: *Clostridium intestinale* PC17, *Anaerostipes* sp. PC18, *Parabacteroides distasonis* PC19 – anaerobic at 37°C, BHI supplemented with hemin (100 mg/L) and vitamin K (0.5 mg/L) (BD), *Lactobacillus johnsonii* PC38, *Lactobacillus murinus* PC39 – anaerobic at 37°C, de Man Rogosa Sharpe (MRS) media (Sigma, MO, USA), *Staphylococcus sciuri* PC04, *Kosakonia cowanii* PC08, *Enterococcus faecalis* PC15, *Staphylococcus xylosus* PC20 – ambient air at 37°C, BHI. Germfree C57BL/6J mice were inoculated with 100 μl of a pooled sample of 1×10^9^ colony forming units (CFU) of each isolate in rPBS to generate the PedsCom gnotobiotic line. To generate the CMCom gnotobiotic line, whole cecal contents of a female SPF NOD/Eα16 mouse was collected anaerobically in rPBS and 100 μl of the slurry was gavaged into germfree C57L/6J mice.

### Sample Collection and DNA isolation

Fecal pellets or intestinal contents were collected in sterile microcentrifuge tubes and stored at −80°C. Bacterial DNA was isolated from samples using the DNAeasy PowerSoil kit (Qiagen, Germany) according to manufacturer instructions.

### 16S rRNA metagenomic sequencing and functional prediction

The V4 variable region of the 16S rRNA gene was sequenced by the PennCHOP microbiome program sequencing core on the Illumina MiSeq platform as previously described^52^. Sequenced libraries were processed using QIIME2 (ver. 2020.6)^53^ and de noised and clustered into amplicon sequence variants (ASVs) with Deblur^54^. Taxonomic classification was performed with SILVA 99% rRNA reference database (ver. 138). Taxa bar plots, alpha diversity and beta-diversity analysis were performed with the R package phyloseq^55^. 16S rRNA metagenomic sequence based predicted metabolic function was conducted using the Tax4fun R package^56^.

### Whole genome sequencing

Bacterial genomic DNA was isolated using the Blood and Cell Culture DNA kit and 500/G genomic tips (Qiagen) according to manufacturer instructions. DNA purity was assessed with NanoDrop 2000 UV-Vis Spectrophotometer (ThermoFisher, MA, USA) and long-read sequencing was performed using the RBK-004 rapid barcoding kit and minION sequencer (Oxford Nanopore Technologies, UK) according to manufacturer’s instructions. Short reads were sequenced by the PennCHOP microbiome program sequencing core using the Nextera XT library preparation kit (Illumina, CA, USA) and the Illumina HiSeq 2500. Contaminating reads were identified using Mash screen^57^. Long and short read sequences were assembled using the unicycler pipeline^27^. Assembled genomes were annotated using the RAST tool kit (RASTtk) on the Pathosystems Resource Integration Center (PATRIC) webserver^28,29^. Average nucleotide identity (ANI) was calculated using the ANI Calculator program with the OrthoANIu algorithm^58^.

### Predicted metabolic function

Assembled genome files of ASF and Oligo-MM12 isolates were obtained from PATRIC. Tables of KEGG orthologs (KO) for each consortium were generated by genome annotation through the KEGG automatic annotation server (KAAS)^30^. Analyses of predicted metabolic function were conducted using the MicrobiomeAnalyst R package^59^.

### Flow cytometry sample staining and acquisition

Mouse lymphoid cell populations of the small intestine, cecum, and colon lamina propria were prepared as previously described^60^. Splenocytes were processed in parallel and used as internal processing and staining controls. Peyer’s patches from the small intestine and the cecal patch was removed before these tissues were used for the preparation of the lamina propria cell suspensions. Stained cell populations were analyzed on the LSRFortessa (BD) and data analysis was performed on FlowJo v10 software (BD). Fluorophore-conjugated antibodies used in staining panel: BV510-CD45 (clone 30F11; Biolegend, CA, USA), PE-Cy7-TCRβ (clone H57-597; Biolegend), APC-Cy7-CD19 (clone 6D5; Biolegend), AF700-CD8 (clone 53-6.7; Biolegend), FITC-CD4 (clone Rm4-5; Biolegend), APC-Foxp3 (FJK-16s; eBioscience, MA, USA), PE-Rory (clone B2D; eBioscience), Pacific Blue-Helios (clone 22F6; Biolegend).

### Reverse transcription quantitative real-time PCR

Two-millimeter tissue sections were excised 5 cm from the distal end of the small intestine (ileum) and stored in RNAlater (ThermoFisher) overnight at 4°C, then at −80 °C for long-term storage. RNA was extracted from tissue samples using the E.Z.N.A. total RNA kit I (Omega Bio-tek, GA, USA) according to manufacturer instructions. cDNA synthesis was performed using the High-Capacity RNA-to-cDNA Kit (Thermofisher) with 2 μg of input RNA. Data acquisition was performed on the 7500 Fast Real-time PCR system using SYBR Green PCR Master Mix (Thermofisher). Cycle threshold (Ct) values of *Tnfa* were normalized to mean Ct values of *Gapdh* endogenous controls. Data analysis was performed using the 7500 software v2.3 (Thermofisher). The 2^-ΔΔCt^ method was used to analyze the relative fold-change of *Tnfa* at 14 and 28-days-old, with baseline levels set at 14-days old in each community. Murine *Tnfa* and *Gapdh* primer pairs were obtained from the PrimerBank database^61^.

### Serum ELISA analysis

Euthanized mice were bled orbitally, and the serum was isolated by a 15 min incubation of blood at room temperature followed by centrifugation at 587 x g for 15 minutes at 4°C. Isolated serum was aliquoted and stored at −20°C. Serum ELISA analysis was performed according to standard protocols. Briefly, 96-well MaxiSorp Immuno plates (ThermoFisher) were coated with unconjugated capture polyclonal antibody goat anti-mouse IgA, IgG1, IgG2b, IgG2c, IgG3, IgM, IgE (Bethyl laboratories, MA, USA) in coating buffer (2.93 g/L NaHCO_3_, 1.59 g/L Na2CO_3_, pH 9.6) overnight at 4°C. Coated plates are washed with wash buffer (0.1% Tween-20 PBS) and incubated at room temperature with blocking buffer (2% BSA PBS) for one hour. Serum samples are diluted in blocking buffer and incubated on plates for 1.5 hours. Plates are washed and HRP-conjugated detection antibodies are incubated with the plates at room temperature for one hour. Plates are developed with the OptEIA TMB substrate reagent set (BD).

### Microbial flow cytometry

Fecal pellets were homogenized in 1 mL PBS, 1% BSA and centrifuged at 100 x g for 15 minutes to pellet solid material. Fecal supernatants are transferred to sterile microcentrifuge tubes and centrifuged at 8000 x g for five minutes to pellet the bacterial fraction and resuspended to an OD600 of 0.1 in PBS, 1% BSA. Bacteria were fixed with 4% paraformaldehyde for 20 minutes at room temperature. Bacteria were washed and resuspended in blocking buffer (PBS, 1% BSA, 20% normal rat serum) and incubated for 20 minutes at 4°C. To determine the precent of fecal microbes bound by secreted IgA, the processed fecal samples were incubated with PE-conjugated anti-mouse IgA antibodies (clone mA-6E1; eBioscience) diluted 1:25 in PBS 1% BSA for 20 minutes at 4°C, washed and incubated in SytoBC nucleic acid stain (ThermoFisher) diluted 1:500 in TBS, for 15 minutes at room temperature. To determine the percentage of fecal microbes bound by systemic immunoglobulins, the processed fecal samples are incubated with autologous diluted serum and stained as previously described with the following fluorophore-conjugated antibodies: goat anti-mouse PE-IgA and PE-Cy7-IgG1 (clone RMG1-1; Biolegend). Flow cytometry analysis on stained bacteria populations was performed on the LSRFortessa and data analysis was performed on FlowJo ver. 10 software. SytoBC and isotype-stained controls were used to define the gates for Ig-coated microbes.

### Salmonella infection model

A streptomycin resistant mutant of *Salmonella typhimurium* SL1344 was cultured with Luria-Bertani (LB) media in ambient air at 37°C. Cultures of *S. typhimurium* were resuspended in sterile PBS and 10–17-week-old mice were infected with 5×10^8^ CFU by oral gavage. *Salmonella* colonization was enumerated 24 hours post infection (h.p.i) by serial dilution of fecal homogenates on LB, 100 μg/ml streptomycin. Cohorts of infected mice were tracked longitudinally for mortality rates. A separate cohort was sacrificed 36 h.p.i. and the Peyer’s patches, spleen and liver were collected for *S. typhimurium* enumeration as previously described. Small intestine and large intestine tissue samples were collected for histopathological assessment.

### Histopathological Assessment

Small intestine and large intestine tissue samples from *S. typhimurium* infected mice and healthy controls were fixed in 10% phosphate-buffered formalin for >24 hours and routinely processed and embedded in paraffin blocks (FFPE). FFPE tissue blocks were routinely sectioned and stained with hematoxylin and eosin (H&E) on glass slides. Histopathological examination was performed blinded to experimental groups by a veterinary pathologist using light microscopy. Samples of small intestine and large intestine were evaluated unpaired as an additional layer of blind evaluation. Detailed assessment of lesions included descriptive evaluation of tissues with morphologic diagnoses as well as semiquantitative scoring of the degrees of inflammation and associated findings, including degree of edema, crypt hyperplasia, goblet cell loss, and frequency of crypt abscesses in each tissue. Scores for each parameter were assigned from 0 (unremarkable) to 4 (severe) at half point intervals.

### Statistical Analysis

Log-rank Mantel-Cox tests were performed with Prism 8 (GraphPad Software Inc). RT-qPCR statistical analysis was performed using independent sample t-tests on mean values with the ggpubr package. All other statistical analyses were performed using non-parametric Mann-Whitney-Wilcoxon tests on median values by either base R or the ggpubr package. P values <0.05 were considered statistically significant. Outliers were determined by z-score analysis.

## Resource Availability

Further information and request for resources and reagents should be directed to and will be fulfilled by the lead contact, Michael Silverman

## Materials Availability

The members of the PedsCom consortium are available upon request.

## Data and Code Availability

All Code will be made publicly available using standard repositories such as github.

